# Across (conformational) space and (relaxation) time: using coarse-grain simulations to probe the intra- and interdomain dynamics of the tau protein

**DOI:** 10.1101/2025.07.21.665865

**Authors:** Jules Marien, Chantal Prévost, Sophie Sacquin-Mora

## Abstract

The biological importance of intrinsically disordered proteins (IDP) has been established for over two decades, yet these systems remain difficult to characterize, as they are better described by conformational ensembles instead of a single reference structure for their folded counterparts. Tau is a prominent member of the IDP family which sees its cellular function regulated by multiple phosphorylations sites, and whose hyperphosphorylation is involved in neurodegenerative diseases such as Alzheimer’s. We use coarse-grain MD simulations with the CALVADOS model to investigate the conformational landscape of tau without and with phosphorylations. Characterizing the local compactness of IDPs allows us to highlight how disorder comes in various flavors, as we can define different domains along the tau sequence. We define the IDP’s Statistical Tertiary Organization (STO) as the average spatial arrangements of domains, which constitutes an extension of the tertiary structure of folded proteins. We also use IDP specific metrics to characterize the local curvature and flexibility of tau. Comparing the local flexibilities with T2 relaxation times from NMR experiments, we show how this metric is related to the protein dynamics. A curvature and flexibility pattern in the repeat domains can also be connected to tau binding properties, without having to explicitly model the protein’s interaction partner. Finally, we rediscuss the original paperclip model that describes the spatial organization of tau, and how phosphorylations impact it. The resulting changes in the protein intra- and interdomain interaction pattern allow us to propose experimental setups to test our hypothesis.

**TOC graphic:** 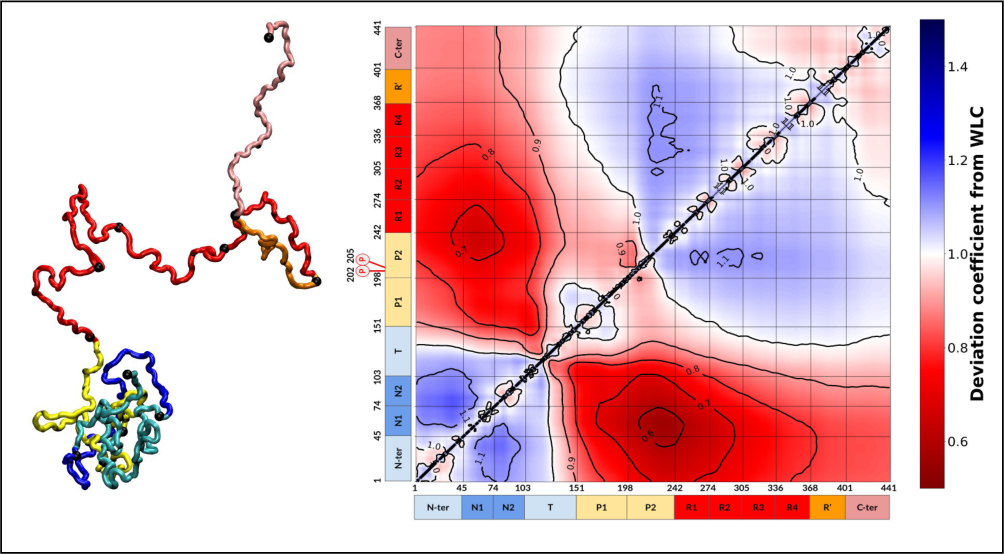

## INTRODUCTION

After over half a century focusing on folded proteins structure,^1^ which culminated with the 2024 Nobel Prize for Chemistry being attributed to AlphaFold,^2^ the last decades have seen a growing shift from the classical structure-function paradigm, ^3^ as the scientific community acknowledged the fact that a large fraction of protein sequences does not fold in isolation while still being functional.^4–6^ Current estimates indicate that roughly 30-40% of eukaryotic proteomes contain intrinsically disordered proteins (IDPs) or folded proteins presenting an instrinsically disordered region (IDR),^7,8^ a proportion that reaches up to 58% of the human proteome.^9^ IDPs and IDRs play a central part in numerous cellular processes,^10,11^ such as transcription, signaling, ^12,13^ transport or cellular organization via phase separation. ^14,15^ Protein disorder is also involved in a wide range of diseases, in particular neurodegenerative disorders,^16–18^ which makes IDPs a lead target for drug development.^19^

Tau is a microtubule-associated protein (MAP) and a well known member of the IDP family, as it plays a part in controlling microtubules (MTs) assembly and the motility of motor protein along MTs.^20–23^ In healthy cells, its function is regulated by reversible phosphorylation.^24^ However, tau hyperphosphorylation has been shown to decrease the protein’s affinity for MTs, to impair its function in tubulin assembly and to induce tau aggregation, which in turn will lead to the formation of neurofibrillary tangles involved in several pathologies (tauopathies) ^25–27^ such as Alzheimer’s disease (AD).^28–30^ More generally, IDPs are a common target for phosphorylations, with important consequences on their biological activity.^31,32^

Due to their conformational heterogeneity, the characterization of IDPs cannot rely on a single reference structure. These systems are better described through conformational ensembles (CoEs)^33–36^ that can be produced with computational tools, such as molecular dynamics (MD) simulations,^37^ which will sometimes integrate constraints derived from experimental data (like NMR or SAXS).^38^ However, simulating long IDPs on the atomic scale remains computationally demanding due to their rugged conformational space and their large fluctuations in size, which is why coarse-grained models offer promising perspectives for CoE generation at low computational cost compared to all-atom simulations. ^39,40^ In this paper, we used the CALVADOS^41^ coarse-grain model to investigate the internal dynamics of full-length tau (which comprises over 400 residue), first without phosphorylations, and then when phosphorylated on Ser202 and Thr205, two sites from the AT8 epitope which are relevant for AD.^42^

Characterizing the resulting CoEs also requires new metrics, as the common similarity measures used to compare folded protein structures, like the root mean square deviation (RMSD) and fluctuations (RMSF), are no longer relevant in disordered system since they require to identify a reference structure upon which all others will be aligned.^43,44^ Previous works such as the Neq metric by Barnoud et al.^45^ and the writhe metric by Sisk et al.^46^ showed promising results for extracting valuable dynamic information from local considerations of the CoE. In this perspective, we employed a new set of metrics, Local Curvature (LC) and Local Flexibility (LF), that were specifically tailored for disordered systems, ^47–49^ and do not require structural alignment since they are based on internal coordinates. In particular we compared the LF values with the T2 relaxation time from NMR experiments and connected the local dynamics of tau to its biological function.

We probed the general and local compactness of tau and their change upon phosphorylation by looking at the average inter-residue distances and their deviation from values obtained for a homogeneous wormlike chain model. This approach enables us to highlight the local heterogeneity within a fully disordered system. We also assessed the impact of phosphorylations on intra and interdomain distances in tau in the monomeric and dimeric forms. Finally, we used these results to reassess the *paperclip* model proposed by Jeganathan et al.^50^ for the structure of tau, and propose new anchoring points for probes (residues S198-T220 and G60-T220) which would permit to test our hypothesis via Förster resonance energy transfer (FRET) or Paramagnetic Relaxation Enhancement/Interference (PRE/PRI) experiments.

## MATERIAL AND METHODS

### General description of tau

The tau protein is a 441-amino-acid long Intrinsically Disordered Protein (IDP) in its longest isoform 2N4R. ^51^ The definitions of tau domains are not unique and can often differ between studies.^22,51–53^ From N-ter to C-ter, we will thus define the domains as follows: N-ter domain (residues 1-44), N1 domain (residues 45-73), N2 domain (residues 74-102), T domain (residues 103-150), P1 domain (residues 151-197), P2 domain (residues 198-241), R1 domain (residues 242-273), R2 domain (residues 274-304), R3 domain (residues 305-335), R4 domain (residues 336-367), R’ domain (residues 368-400) and C-ter domain (residues 401-441). We also define 4 main regions: the N-ter, N1, N2 and T domains as the projection region; the P1 and P2 domains as the Proline-Rich Region (PRR); the R1-4 and R’ as the repeat domains region and the C-ter domain as the C-ter region. A schematic description can be found on Figure 1. As far as we know, the domain from residues 103 to 150 was never properly named before. We name it the Turn (T) domain for reasons further explained in the discussion. The ability of tau to bind to MTs is principally mediated by its repeat domains and the PRR.^51,54^ Work by Brotzakis et al. further refined our understanding of the binding dynamics by identifying weak and strong binding patches, centered on sequence motifs VQIXXK and SKXGS respectively.^55^

**Figure 1:**
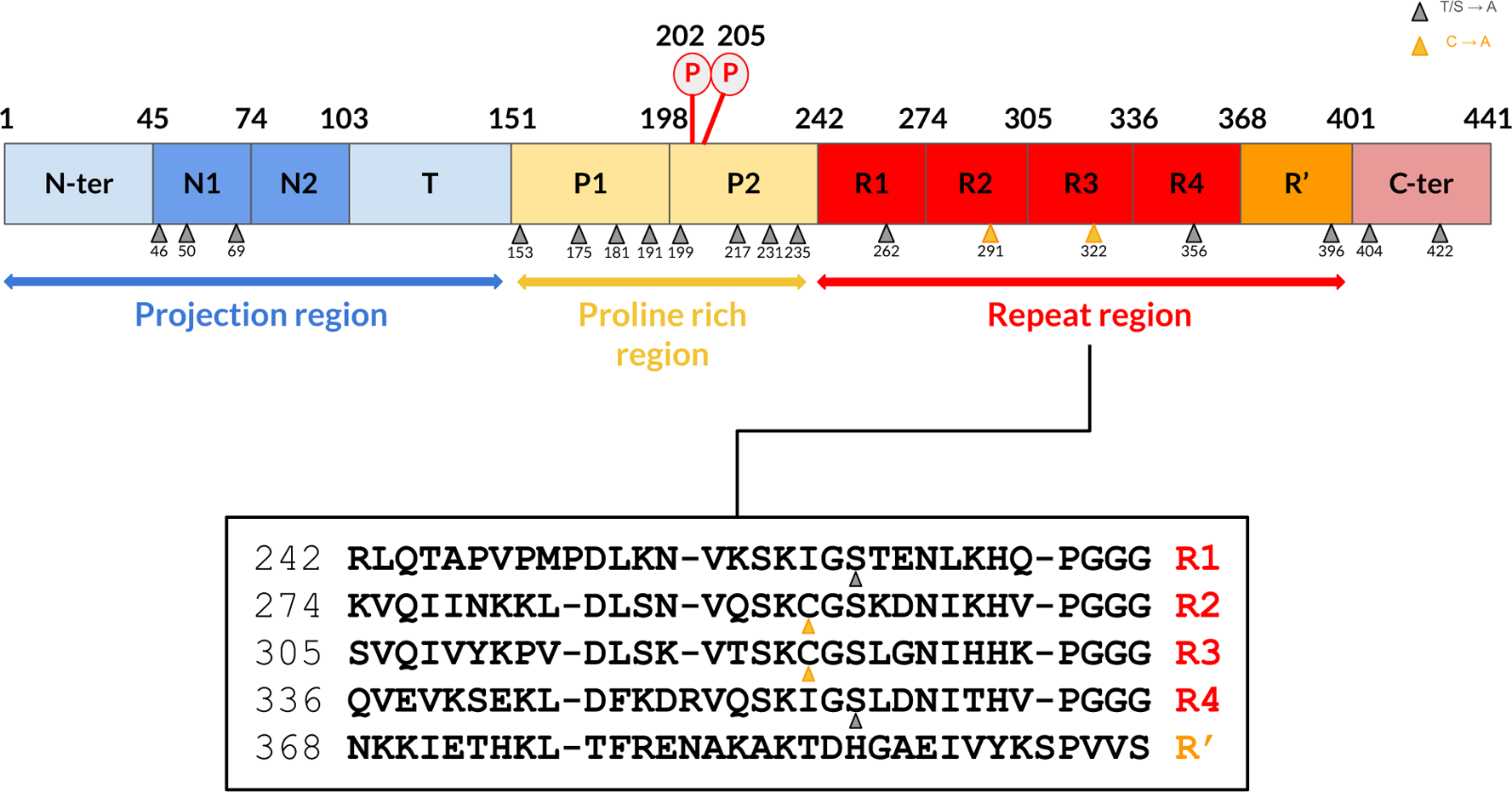
Domains and regions of tau, and alignment of the sequences of the repeat domains. Mutations to alanines are signaled by a grey triangle. Natural cysteins positions are signaled by a yellow triangle. Sequence alignment was performed with the MAFFT algorithm.^56^

Inducing site-specific phosphorylations on a protein, especially an IDP, is not a simple task experimentally. For this reason, Lasorsa et al. decided to engineer synthetic mutants of the tau protein that display alanine mutations at phosphorylation sites.^57^ Furthermore, the natural cysteines of tau were also mutated to alanines and moved at S185C and A227C, closer to the AT8 epitope. The AT8 antibody binds to paired-helical filaments of tau and has a specificity for the pS202/pT205 and pS202/pT205/pS208 phosphorylation states of the protein.^42^ We define the AT8 epitope as the region of tau located between residues 195 and 214 in agreement with the peptide used for its characterization. The summary of the mutations can be found on Figure 1 as grey and yellow triangles. A collaborative study combining experiments and simulations was previously performed to show that the introduction of these mutations does not modify the dynamic behavior of the protein on the global and local scales (see Ref^49^).

### CALVADOS simulations

The sequence of the tau mutant was extracted from the work of Lasorsa et al.^57^ We performed a simulation of the unphosphorylated monomer using a modified version of the IDRLab notebook with the CALVADOS 2 parameters in order to later accommodate for phosphoresidues.^40^ The CALVADOS 2 model represents each residue as a single bead of varying size, charge and hydrophobicity, centered on the C*_α_* and with an implicit consideration of the solvent which allows for highly efficient computations. A second simulation with phosphorylations at Ser202 and Thr205 was run with phosphoparameters extracted from the work of Rauh et al., ^58^ and compared with the pCALVADOS forcefield (Figure SI-1).^49^ Since parameters for phosphoresidues did not exist for CALVADOS until very recently, we had implemented the parameters of a similar model by Perdikari et al., ^59^ which ended up being very similar to those derived by Rauh et al. As no major difference was spotted between the simulation with the latter parameters and pCALVADOS, we adopted Rauh et al. phosphoparameters as they were more extensively tested and specifically designed to work within the CALVADOS distribution.^60^

Simulations were performed with OpenMM v7.7,^61^ with a Langevin integrator of friction coefficient 0.01ps*^−^*^1^ and a 10fs timestep in the NVT ensemble. The temperature was set to 278 K and the salt concentration to 0.08 mol.L*^−^*^1^ to reproduce NMR experiments. Histidines were uncharged while terminal residues were kept charged. Monomeric trajectories underwent a 100 000-step equilibration, followed by 100 000 000 steps of production. These parameters are in agreement with previous successful studies from the conceptors of the CALVADOS forcefield,^40,62^ and we previously showed that these parameters ensured convergence.^49^ 20 000 evenly-spaced frames were collected from the complete production step after discarding the initial equilibration part, and backmapping to all-atom was performed following the same protocol as in Lohberger et al.^49^

Simulations of phosphorylated and unphosphorylated tau mutant dimers were performed using the CALVADOS package.^60^ Two tau mutant monomers were randomly placed in a box of sidelength 20 nm with all other parameters being similar as for the monomeric case, except for the equilibration and production times, which were doubled to ensure convergence. All analyses were performed using the MDAnalysis and MDTraj python packages.^63,64^

### Proteic Menger Curvatures, Local Curvatures and Local Flexibilities

We previously introduced Proteic Menger Curvatures (PMCs) as a way to define Local Curvatures (LCs) and Local Flexibilities (LFs), two metrics which allowed us to characterize the curving and rigidification induced by counterion-mediated bridges in multiphosphorylated peptides.^47^ Briefly, the PMC of a residue can be defined as the inverse of the radius of the circle passing through its *C_α_* and the *C_α_* of its *s*-nearest neighbors, where *s* is defined as the spacing and is usually set to 2 for proteic backbones.^48^ As a result, the PMC of the first and last two residues is not defined. The LC and LF associated to the residue are respectively the average and standard deviation on its PMCs across all sampled conformations. Comparison of LCs and LFs in multiphosphorylated CALVADOS simulations revealed a local elongation and rigidification near the phosphorylation sites, proportional to the number of phosphorylations.^49^ We used the Menger_Curvature MDAKit package to calculate the PMCs, LCs and LFs of tau in all simulations. ^48,65^

### Persistence length and the Worm-Like Chain model

We define the length of an amino acid as *l_a_* = 3.8 Å, as defined in the CALVADOS model.^41^ We can then define the contour length *L* of an IDP with *N* residues as *L* = *N* ∗ *l_a_*. The persistence length *P* of a polymer is a simple measure of its stiffness, and can be shown to follow the relationship:

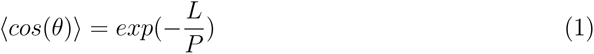

with *θ* the angle between 2 bond vectors. The right-hand side of Eq 1 can be determined by calculating the autocorrelation function between the bond vectors. The calculation of the persistence length of an IDP monomer in our simulations was performed using the **polymer.PersistenceLength()** method in the MDAnalysis package.^63^ Unless specified otherwise, the persistence length *P* of the unphosphorylated monomer simulation was measured to be 6.6 Å, and *P* will be set as such.

A simple approximation to model polymeric chains is to represent them as a Worm-Like Chain (WLC).^66^ It assumes that the polymer is an isotropic rod, flexible but displaying a certain rigidity, which is solely parameterized by an associated persistence length. Highly flexible proteins like IDPs can be treated by this approximation and be considered as polymer chains of *N* amino-acids. It can be shown that the average squared distance 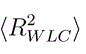 between points *i* and *j* spaced by *ℓ* in a WLC follows the equation:

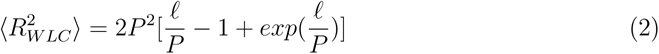

By defining *ℓ* = *n* ∗ *l_a_*with *n* the number of residues between *i* and *j*, and taking the square-root of Eq 2, we can define a deviation coefficient *d_W LC_* from the WLC by taking the ratio with the average distance ⟨*R_sim_*⟩ between two *C_α_* in a simulation:

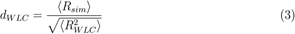

If *d_W LC_ >* 1, the average distance between the two *C_α_* in the simulated IDP is higher compared to the WLC model with a similar persistence length *P* . A similar distance normalization was employed by Holla et al using a polynomial expression for the theoretical inter-residue distance,^67^ and by Lindorff-Larsen et al. using an interresidue contact map generated from the simulation of a random coil. ^68^ Alston et al. recently developed a comparison model called the analytical Flory random coil (AFRC) and showed that the WLC was also a relevant reference to use for end-to-end distances. ^69^

### Secondary chemical shifts

Chemical Shifts (CSs) and T1 and T2 relaxation times were obtained from Lasorsa et al. ^57^ CS determination is widely used to assess the accuracy of a CoE obtained by MD simulations, but CSs suffer from a dependency in amino-acid type. ^70^ One can remove it by calculating Secondary Chemical Shifts (SCSs), defined as the difference between the measured CS and the random-coil CS of the same amino-acid type. We used the random-coil CS values measured by Kjaegaard et al. to calculate our SCSs. ^71^ CS values were predicted with SPARTA+ every 10 frames and averaged over the course of the trajectory.^72^

### Analyses

Sequence analysis of the tau protein was performed using the localCIDER python package.^73^ The SeqDYN webserver^74^ with default parameters was used to predict R2 values for the tau mutant sequence, then inverted into T2 values in order to be compared to experimental and computed T2. Results were displayed with the Matplotlib python package^75^ and snapshots were generated using Visual Molecular Dynamics (VMD).^76^

## RESULTS

### Sequence composition hints at dynamical properties

Relating the sequence composition to the dynamics of IDPs is a complex endeavour, which has seen significant improvements in recent years.^67,77,78^ The complete tau sequence comprises equal fractions (13%) of negatively and positively charged residues (see Table SI-1) and should therefore belong to the *globules* group as defined by Das et al.^78^ However, these charged residues are unevenly distributed along the sequence, and the N-terminus harbors a large number of negative charges.^52^ With 23% of negatively charged residues in the projection region and 17% of positively charged residues in the PRR, one can hypothesize that these regions will tend to attract each other, but how they will do so is a more difficult problem. We calculated the patterning coefficient *κ*, which quantifies the segregation of positive and negative charges along the protein sequence, ^77^ for different parts of the sequence of tau using the localCIDER package.^73^ *κ* is bounded between 0 and 1, with 0 reflecting a complete homogeneity of the charge distribution, while 1 indicates that all positive charges are separated sequentially from the negative charges. The complete tau sequence has a *κ* of 0.18 while the combined sequences of the projection region and the PRR have a *κ* of 0.22. This correlates with their respective charge composition, and implies a possible “hairpin” behavior, since sequences of higher *κ* have a propensity to adopt hairpin-like conformations according to Das et al.^77^ Where the turn is located, however, cannot be determined from the sequence alone. On the other hand, the repeat domains have a *κ* value of 0.10, which would imply a more extended conformation. ^79^ The *κ* coefficient might however be less reliable for proline-rich sequences,^78^ and the size and complex domain composition of the tau protein makes it more difficult to draw detailed conclusions from sequence analysis only.

### Across conformational space…

The CALVADOS forcefield was developed based on global observables such as SAXS profiles and more local observables such as NMR paramagnetic relaxation enhancement data. ^41,62^ CALVADOS simulations of a tau mutant yielded similar hydrodynamic radii for unphosphorylated and multiply phosphorylated forms as those measured with dynamic light scattering experiments.^49^ We therefore assumed that the inter-domain interactions within a monomer modelled with CALVADOS would be accurately described.

### Tau monomer

We first characterized the intramolecular conformational space of the tau mutant by simulating a single monomer. An example of a sampled conformation is displayed on Figure 2. We computed the average distance map for unphosphorylated and ERK2-phosphorylated monomers (Figure 3A). The P1 and P2 domains clearly exhibit a higher compaction than the rest of the sequence, with most of their residues being on average distant by less than 60 Å in contrast to more than 80 Å for residues with a similar sequence spacing in the repeat domains. This higher compaction of the projection and PRR regions compared to the repeat region can be visualized on Figure 2 and is in agreement with FRET measurements performed by McKibben et al. ^54^ The distance isovalue at 60 Å also reveals an expansion of distances between domains P2 and N1 upon phosphorylations at S202 and T205.

**Figure 2:**
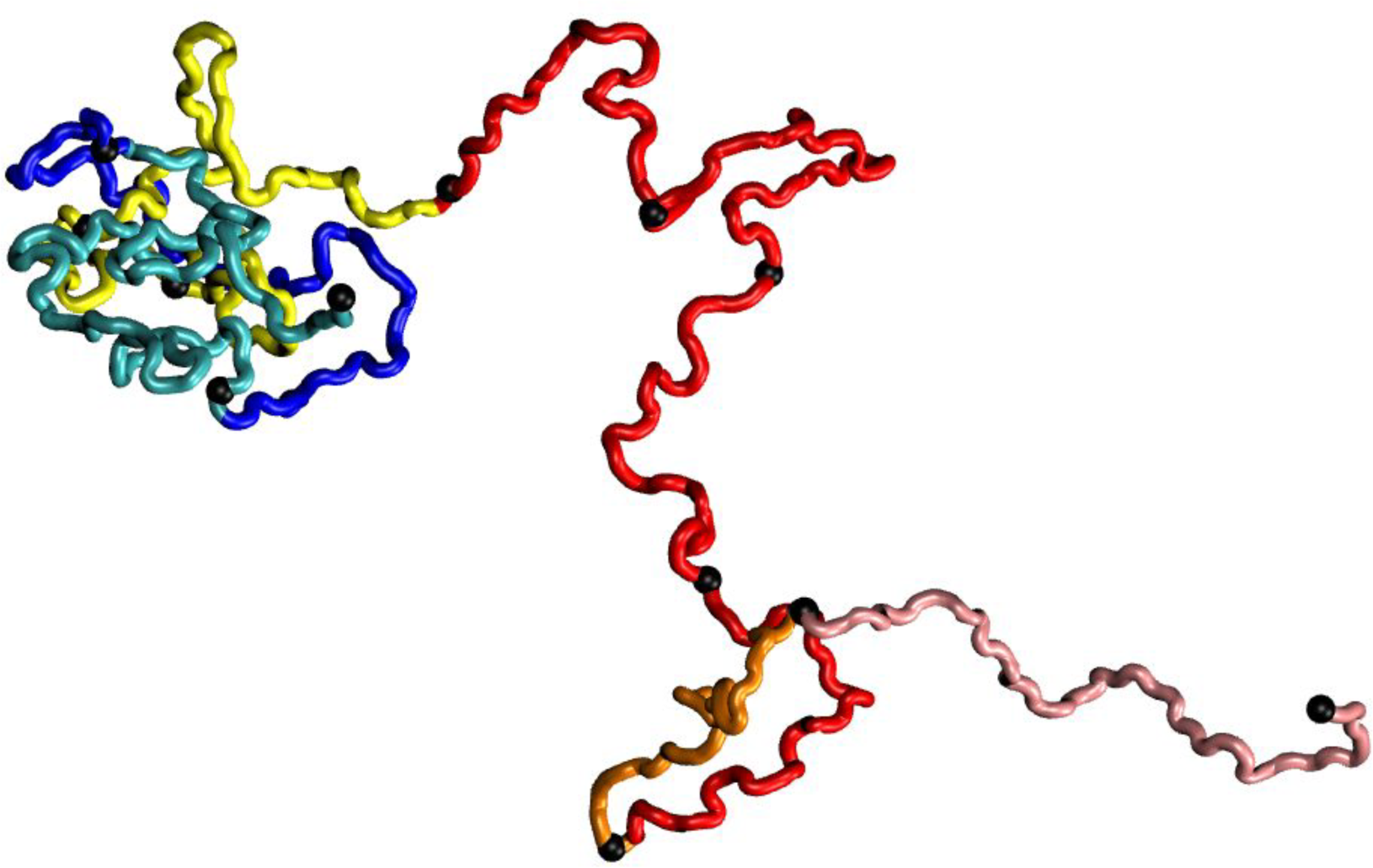
A representative snapshot of the unphosphorylated tau mutant. The color scheme for the tau domains is the same as in Figure 1. The black Van der Waals spheres hightlight the frontiers between the domains defined in Figure 1. Image rendered with Visual Molecular Dynamics.^76^

**Figure 3:**
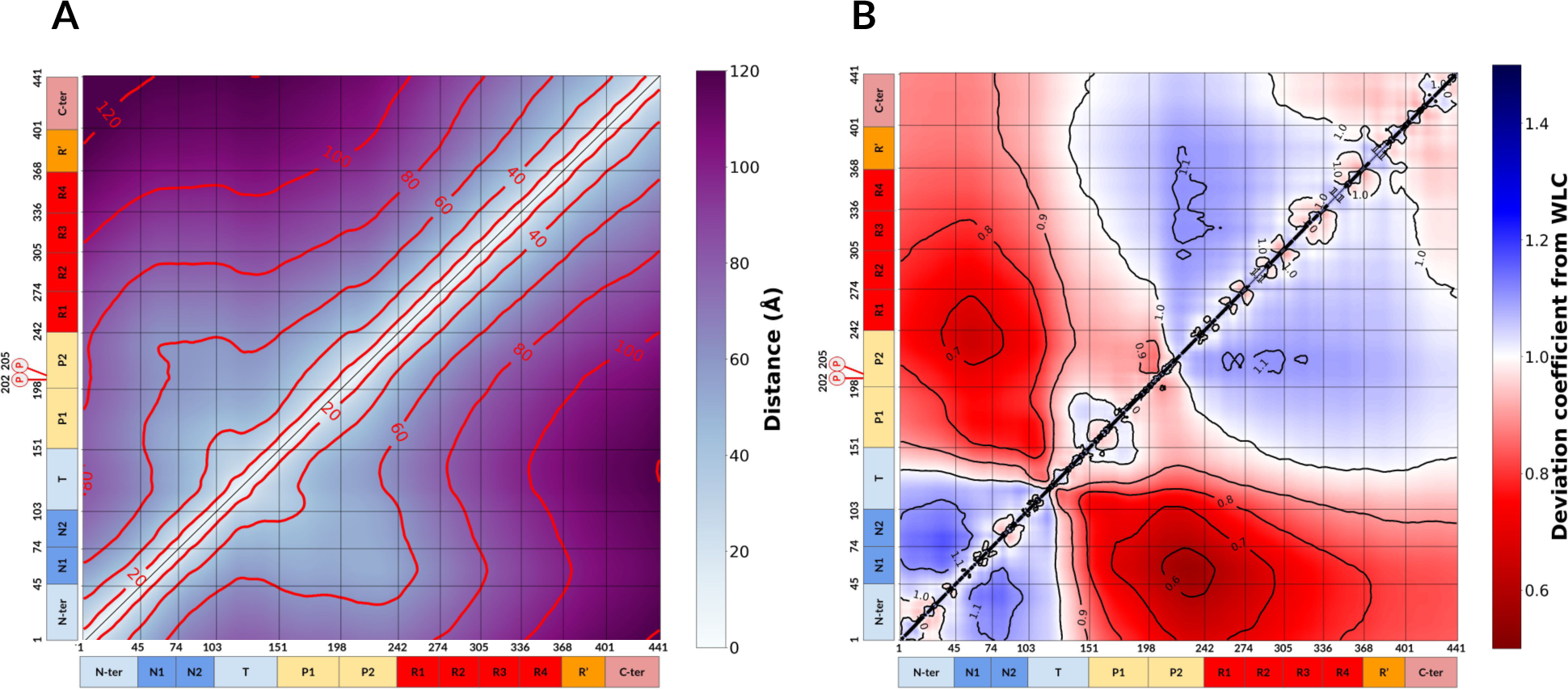
A) Average distances between residues in the unphosphorylated (lower part of the matrix) and ERK2-phosphorylated (upper part of the matrix) tau mutant monomers. Increased distance is visualized by a deepening of the purple color. Isovalues 20Å, 40Å, 60Å, 80Å, 100Å and 120Å are plotted as red isolines. B) Deviation coefficient matrix from the WLC model. Deepening of the red color indicates a shorter average distance compared to the WLC model, deepening of the blue a longer average distance. Isovalues 0.6, 0.7, 0.8, 0.9, 1.0 and 1.1 are plotted as black isolines.

In order to probe the inter-domain dynamic organization and the heterogeneity of tau local compactness, we normalized the average distance map by distances predicted by the WLC model (Figure 3B). The resulting map reveals a clear patterning of the inter-domain spatial organization. The N-ter, N1 and N2 domains are not as close to each other as predicted by the WLC, and the repeat domains as well, meaning that those parts adopt more extended conformations. The C-ter domain is slightly closer than predicted (deviation coefficient of 0.9) to the R’, R4 and R3 domains and to itself, suggesting that its residues bend slightly toward the extended repeats. The most striking deviation from the WLC model is that of the P2/N1 distance. In the unphosphorylated form, these two domains display a deviation coefficient of less than 0.6, indicating an average spatial proximity far greater than prescribed by the WLC. Overall, the N-ter, N1 and N2 domains are closer to the PRR, the repeats and the C-ter domains. The clear delimitation between over- and undercompact distances drawn from the middle of the T domain can be interpreted as a turn in the polymer which will be further discussed below. One can also notice a periodicity in the intra-domain interactions of the repeat domains, suggesting that their respective N-terminal part is more extended than their C-terminal part, in correlation with the *β*-sheet propensities measured by NMR by Mukrash et al.^80^

Phosphorylation induces a lesser compaction of the P2/N1 distance as the deviation coefficient does not reach values below 0.6, and shifts the compaction from residues at the center of the P2 domain to those at the limits with the R1 domain. Except for this difference with the unphosphorylated monomer, the interaction pattern drawn from the deviation coefficient map is not significantly altered, which incidentally confirms that both simulations have converged. The N1/P2 average proximity and the shift in residues induced by phosphorylations can be simply explained by the charged residue distribution in the protein sequence. Indeed, the N1 domain is rich in negatively charged residues (E53,57,58,62,73 and D65) while the P2 domain is rich in positively charged residues (R209,211,230,242 and K214,215,234,240), so it is easy to imagine that these domains would have favorable interactions bringing them closer. When phosphorylations are introduced at S202 and T205, N1 gets repelled from the C-terminal part of P2, inducing the observed shift in the deviation coefficients. This hypothesis is supported by the observation of a 0.9 isovalue in the intra-domain portion of the phosphorylated map: this translates the effect of the negatively charged phosphorylations which “pull” the positively charged sequence of P2 towards them. This representation of the inter-domain organization by comparison with the WLC model is therefore successful in describing overall monomeric dynamics and also subtle changes in interactions induced by Post-Translational Modifications (PTMs) such as phosphorylations.

### Tau dimer

In a second step, we investigated the impact of intermolecular interactions on the interdomain dynamics. At an experimental concentration of 100 µmol.L*^−^*^1^, a dimeric simulation would require a cubic box of length 321 nm. With an average radius of gyration Rg of 5 nm, this would make intermolecular interactions extremely rare. We therefore performed simulations of two tau mutant monomers in a box of length 20 nm. This length allows both monomers to extend at their average Rg while maximizing their interactions. We computed the average distance maps and the deviation coefficient map from the WLC model (Figure SI-2), as well as the intermonomer contact map (Figure SI-3). A contact was defined as two intermonomeric beads being closer than 8 Å.

The overall compaction of the projection region is still present in the dimeric simulation, but the local elongation induced by the phosphorylations is not noticeable anymore on the 60 Å isoline (Figure SI-2A). Modification of the deviation coefficient map with respect to the monomeric case involves a decrease of the distances between the projection regions and the C-terminal domain (Figure SI-2B). Domains N1 and P2 are further away on average as we cannot find the 0.6 isoline in the unphosphorylated case anymore. The N-ter and C-ter domains also get closer to one another as the deviation coefficients decrease from 0.9 to 0.7. The rest of the interaction pattern remains the same for the repeat domains and for the interactions within the N-ter, N1 and N2 domains. Addition of phosphorylations still displaces the 0.7 isoline towards the C-terminal of the P2 domain, and the 0.9 isoline still appears in the intra-domain interactions of P2.

In order to assess whether intermonomer interactions could explain the differences observed in the intramonomer maps between the monomeric and dimeric simulations, we computed the intermonomer contact maps for the dimeric simulations with and without phosphorylations (Figure SI-3). Contacts are scarce, with maxima at 0.2% of the simulation time (which corresponds to around 80 cumulated frames per residue pair). These maxima are however interesting since they occur between residues located at the centers of the N1 domain and the P2 domain. Most of the contacts occur between the N-terminal portion and the PRR, which is in agreement with the repartition of charged residues in these regions. This would probably induce some steric hindrance for the intramolecular interactions of the P2 domains with the rest of the monomer, explaining why the profiles are not exactly the same for the monomeric maps. Phosphorylations displace the intermonomer contacts in the same way that they do for the intramonomeric ones. Contacts between the AT8 epitope and the N-terminal regions are also completely negated upon phosphorylation. Finally, these PTMs also lead to a contact increase between the N-terminal region of the tau mutant and the N-terminal part of the R’ repeat domain, and transient interactions appear between the phosphate groupes and the repeat domains.

These inter-domain characterizations thus provide powerful insights on the dynamics of the tau protein and on the impact of phosphorylations at the AT8 epitope.

## .. . and relaxation time

We next probed the tau intra-domain dynamics by using the Local Flexibility (LF) metrics.^48^ Since the transversal relaxation time T2 is sometimes interpreted as a measurement of backbone flexibility of IDPs,^57^ we first wanted to assess whether a correlation existed between T2 and LFs.

We superposed the LFs results and the T2 values (see Figure SI-4) and calculated the Pearson’s correlation coefficient *r*. For all residues for which a T2 value was measured experimentally, the correlation is mild (*r* = 0.37) even if the trend of LFs looks very similar to that of T2. We also calculated the correlations without the first and last 10 residues, and only for the repeat domains, yielding much mire substatial values of *r* = 0.47 and *r* = 0.59 respectively. A first explanation for the mild correlation on the overall sequence is that the T2 measurements tend to skyrocket for the terminal residues while LFs remain stable in amplitude. We will therefore exclude the first and last ten residues from our analysis from this point on. We also notice clear oscillations at the residue level in LFs which might hide the correlation of the trend.

We decided to treat these oscillations as a noise in the LF signal and to remove this noise with a Savitzky-Golay filter^81^ by finding the optimal parameters (window length and polynomial order) which maximizes the correlation to T2 (Figure SI-5). We used the **scipy.signal.savgol_filter** function from Scipy. ^82^ We selected a window length of 15 residues and a polynomial order of 4. The resulting filtered LFs have a correlation with T2 of *r* = 0.47 on the entire sequence, *r* = 0.59 without the first and last 10 residues, and *r* = 0.67 for the repeat domains, indicating that their trend is indeed correlated. We then performed a linear fit based on the T2 values on the smoothed LFs to convert them into LF-derived T2 values (Figure 4). It should be noted that the goal here is not to create an accurate predictor but to allow for a proper comparison. LF-derived T2 should be treated as a variable possibly correlated to experimental T2 rather than as a definite prediction. Limitations are described in the Supplementary Information.

**Figure 4:**
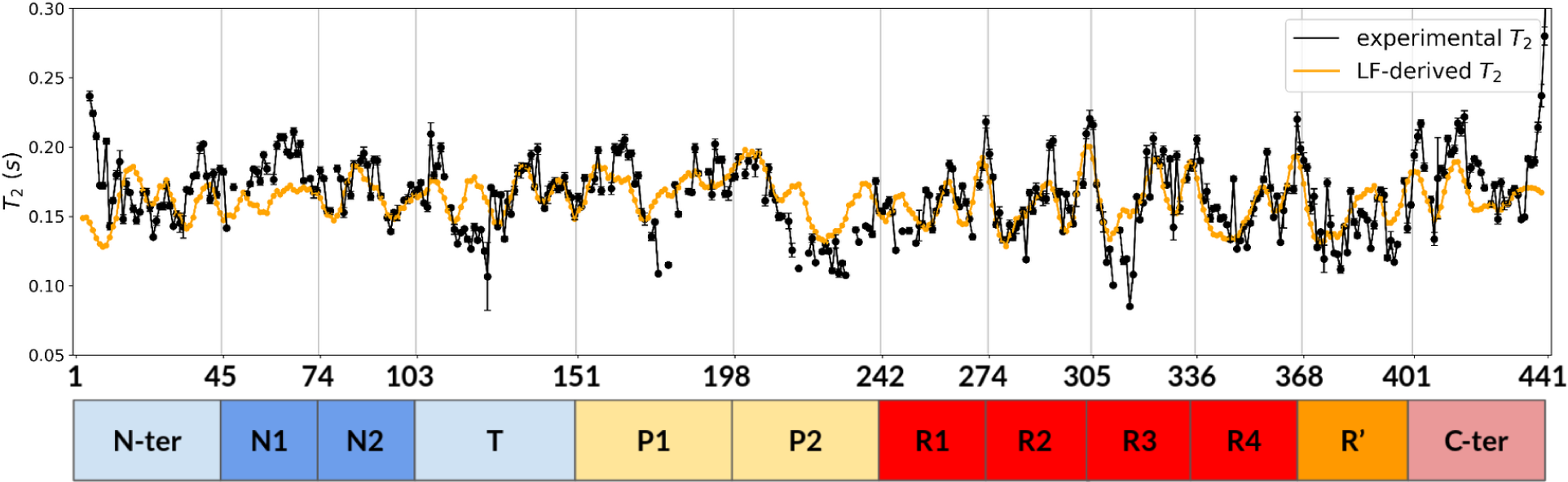
Plot of the experimental T2 values (in black) and the LF-derived T2 values derived from the smoothed LFs of the unphosphorylated mutant (in orange).

The resulting Root-Mean-Squared Error (RMSE) is 0.02 s, which is less than 10% of the average amplitude of the T2 measurements. The trend of the signal is also well reproduced, with a notable exception in the middle of the T domain between residues 115 and 125 and in some parts of the PRR, which are further discussed in the Supplementaries. Since LFs are continuous and smoothed, we were able to notice a pattern in the repeat domains, which we decided to investigate further. We therefore superposed the T2, smoothed LFs and LCs from the repeat domains (Figure 5).

**Figure 5:**
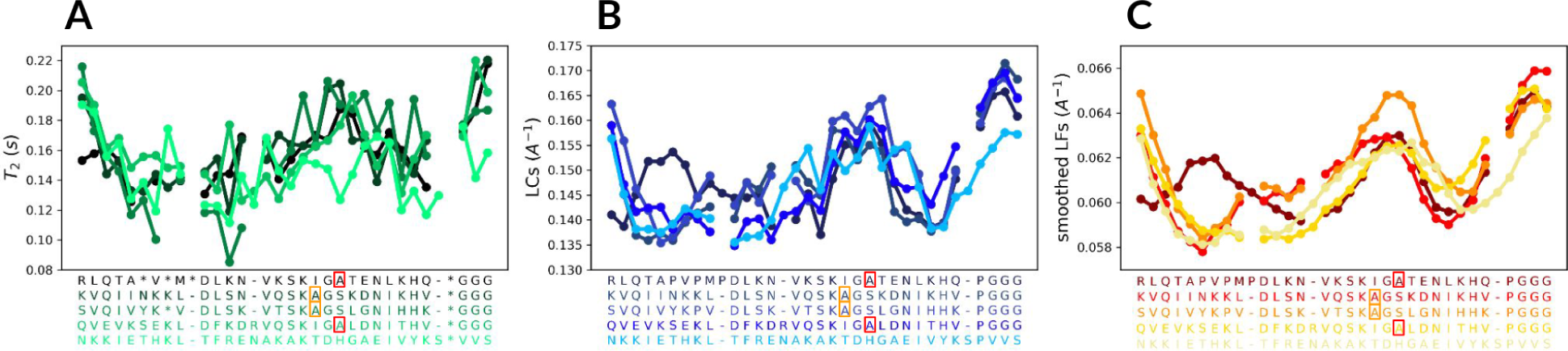
Superposition of the values for the repeat domains (with the sequences underneath the curves corresponding to R1, R2, R3, R4 and R’, from top to bottom / dark to light) for A) T2 experimental values, B) Local Curvatures (LCs) and C) smoothed Local Flexibilities (LFs). Alanine mutations are squared in red, alanine mutations of the natural cysteins are squared in yellow.

The alignement reveals a clear pattern of flexibility in the repeat domains. The C-terminal parts are less flexible than the SKXGS motifs. The four residues neighboring the histidine residues in R1-4 are also less flexible, while the PGGG/PVVS motifs are the most flexible of the sequences. These findings are in full agreement with the study of Stelzl et al., which combined different computational and experimental methods to probe the local structure of the K18 tau fragment (R1-4),^83^ and with the early work by Mukrasch et al., which characterized through NMR the local dynamics of the K32 tau fragment (P2+R1-4+R’, S198-T394). ^80^

The R1 repeat domain displays a higher flexibility and curvature next to its N-terminus than the rest of the repeats, however this is not observed in the T2 signal and constitutes a discrepancy between the LF-derived T2 profile and the experimental T2 (Figure 4). Importantly, the R’ pseudo-repeat behaves very similarly to the other repeats despite their differences in sequence. From the PMC calculations, we can obtain LFs but also LCs, which monitor the average curvature of a residue. The LC profiles of the repeats exhibit a pattern similar to the LFs, as more flexible residues are also more curved in average. This implies that the N-terminal part of the repeats is relatively extended and rigid compared to the SKXG motifs, and that PGGG/PVVS motifs could act as a sort of flexible hinges between the different repeats, in accordance with what was already determined with the deviation coefficient maps (Figure 3).

We also used the smoothing of the oscillations and the linear function to transform the LFs of the phosphorylated monomer into LF-derived T2 values (Figure 6). Similar parameters as for the unphosphorylated case were employed for the smoothing and the linear fit. No major difference could be observed between the LF-derived T2 values of the unphosphorylated and phosphorylated monomers, except at the phosphorylation sites where the flexibility is reduced, similarly to what was observed in a previous study on different phosphorylation states. ^49^ The comparison to experimental data is less accurate than for the non-phosphorylated case with a RMSE of 0.027s. It is also less accurate for the P2 domain, especially near the phosphorylated epitope. This is probably due to the localized strong charges carried by the phosphorylations, which might distort the signal.

**Figure 6:**
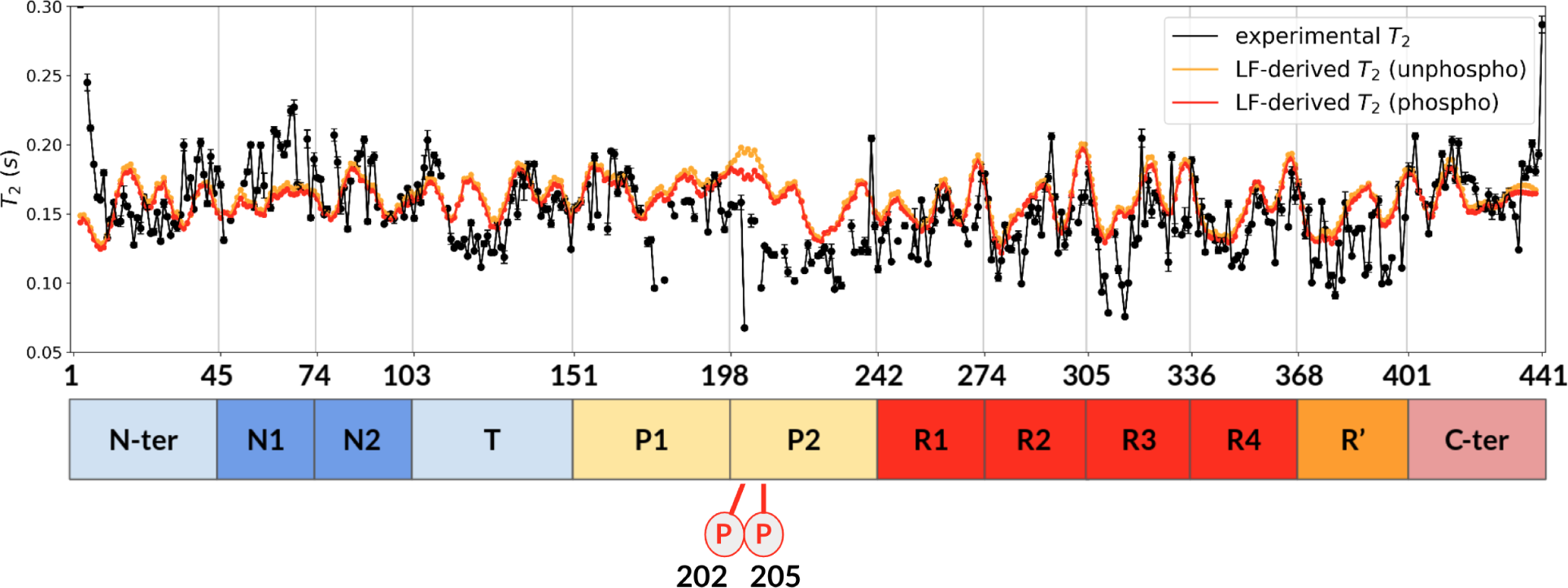
Plot of the experimental T2 values (in black) and the LF-derived T2 values derived from the smoothed LFs of the unphosphorylated (in orange) and phosphorylated mutants (in red).

In order to probe how deeply connected to the dynamics LFs really are, we also compared the unphosphorylated LF-derived T2s to the T2 predictions made with the SeqDYN method by Qin and Zhou (Figure SI-6).^74^ SeqDYN is a machine-learned method which predicts the relaxation rate R2 values from the proteic sequence alone, R2 being the inverse of T2. T2 values are influenced by the strength of the magnetic field used to make the measurements, which explains why the initial prediction values made by SeqDYN are twice as large as the experimental T2. As mentioned by the authors, one can however apply a uniform scaling factor on the results and we determined this factor to be 0.54. The scaled SeqDYN predictions correlate less with the experimental data than the ones derived with smoothed LFs (*r* = 0.54 vs *r* = 0.59) and the RMSE is slightly higher (0.023 s vs 0.020 s). Considering only the repeats amplifies the differences between SeqDyn and LFs based predictions, with the first ones correlating only at *r* = 0.44 with the experimental data (RMSE of 0.026 s), and LFs-derived T2s correlating at *r* = 0.67 (RMSE of 0.019 s) with the experimental data. LCs and LFs only describe properties of the protein backbone, with no direct consideration for the side chains or the specific chemical environment of the residues. We also tried reproducing SCSs to assess whether relevant information could be found at such a small scale from the CALVADOS simulations. The one-bead-per-residue representation does not allow for SCS calculations, the monomer simulations were thus backmapped to all-atom resolution using the scheme described in Ref.^49^ Calculation of SCSs is described in the method section.

As expected, the computed SCSs values for the backbone amide (H_N_) protons do not correlate at all with the experimental measurements (*r* = 0.075 and RMSE of 0.23 ppm) since they are very sensitive to the local chemical environment, which is apparently not well rendered by the backmapping process (see Figure 7). This can easily be understood by considering how CALVADOS represents IDPs. Indeed, the one-bead-by-residue model cannot account for secondary structure propensities, which are crucial for the reproduction of SCSs. The computed SCSs values for the nitrogens themselves on the other hand display a correlation of *r* = 0.62. There is however a general shift towards lower ppm values, and the RMSE reaches 2.044 ppm. These results, combined with the correlation of the LF-derived T2 to the experimental data, indicate that the resolution of the CALVADOS representation limits the level of accurate information that the model can provide to the intra-domain scale, and that the backmapping process is not advanced enough to correctly reproduce the local chemical environment within residues.

**Figure 7:**
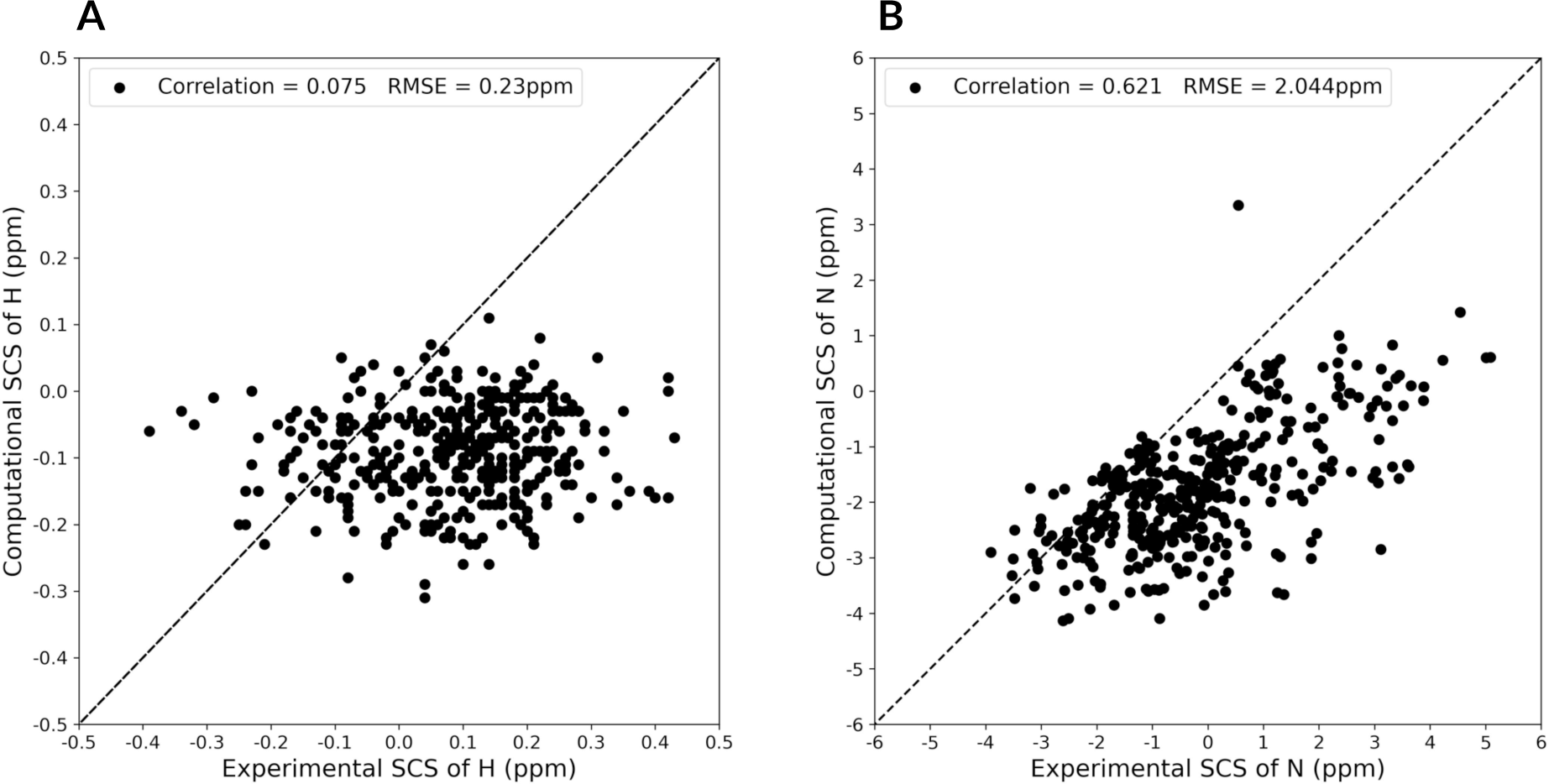
Comparison of the experimental Secondary Chemical Shifts (SCSs) and the SCSs derived from SPARTA+.^72^ A) backbone amide (H_N_) protons, B) nitrogens.

## DISCUSSION

### Overall organization of IDPs

The first noticeable result from this study is that, thanks to the recent experimentally-informed development of coarse grained force-fields such as CALVADOS, it is possible to get precise information regarding the spatial organization of very long IDPs that takes into account their sequence inhomogeneity. Our analyses point to a statistical spatial domain distribution, which we term Statistical Tertiary Organization (STO), and which can be seen as an extension of the tertiary structure of folded proteins. This STO can be physically interpreted. Briefly, certain regions of the IDP show higher probability to be spatially close or far from each other. We must however insist on the idea that the STO is not a conformation, but a resultant of the average behavior of the IDP. Our simulations confirm experimental observations about the average proximity of regions that may be far away in sequence, and the analyses add precise characterization of the sequence-defined domains and their dynamic behavior. Of note is the observed sensitivity of the STO to point mutations or post-translational modifications, that we show may be influenced by long-range interactions within the IDP or between IDP monomers.

We also explore in this work the link between the overall organization of the IDP and the backbone dynamics viewed at the local level. Below, we discuss the possibilities and limitations of the metrics we developed to this aim.

### LFs finely describe backbone dynamics

The high correlation for the tau mutant between the smoothed LFs and the experimental T2 suggests that LFs as a metric is efficiently capturing an important component of the dynamics of the IDP backbone. A visual inspection of the curves comparing the LFs-derived T2 values to the SeqDYN prediction allows to notice that some of the variations of the experimental signal are captured by LF-derived T2s but not by SeqDYN (Figure SI-6). This raises the question of whether some dynamic effects cannot be directly derived from local consideration on the sequence on which the SeqDYN model was based, but can effectively be captured by the LFs. Another useful consideration is the calculation method of LFs: the LF of a residue is the standard deviation on its PMC distribution. Not only do we have access to this distribution and its component values, but they can be easily interpreted structurally. This correlation between the experimentally measured T2s and the computationally derived LFs could therefore lead to a deeper understanding of IDP conformational ensembles and the relationship between local averaged properties and backbone dynamics.

### LFs as a T2 proxy

Deriving T2 values from a MD trajectory is a theoretically advanced task which relies on heavy calculations. We refer the reader to ref^84^ for more in-depth details. Briefly, the most common methodology consists in deriving the spectral density *J* (*ω*) as it can be related to R2. The determination of *J* (*ω*) from the MD trajectory is made by calculating the time correlation function for each bond vector of interest. Recent improvements such as the ABSURD method by Salvi et al.^85^ and later the ABSURDer method by Kümmerer et al. ^86^ yield qualitative and quantitative agreements with NMR data.

In addition to the complex post-processing they require, these methods also need all-atom MD ensembles with a very thorough sampling in order to be able to satisfyingly calculate the spectral density. On the other hand, converged CALVADOS simulations of a single monomer can be achieved within a few hours on a modern laptop, and using the Menger_Curvature MDAKit implementation, the LFs can be calculated in a matter of minutes. The conversion to LF-derived T2 values then requires a simple filtering and a basic linear fit which are computationally inexpensive and easy to interpret. While the accuracy of the prediction is not on par with other state-of-the-art methods, LFs might still be able to provide a fast first estimate of the local flexibility of an IDP that could prove useful, for example for the analysis of large datasets. Further tests on other IDPs should however be performed in order to assess the limitations of the correlation between LFs and T2.

Indeed, we noted some local discrepancies in the LFs-derived T2 even for the unphosphorylated monomer (Figure 4, Figure SI-7). These discrepancies could be exploited to identify new directions to upgrade the CALVADOS description of IDPs (see discussion in SI). In another hand, previous work by Lindorff-Larsen et al. on a 200 µs simulation of an unfolded protein showed significant correlation between transient contacts and J(0), an observable related to T2.^87^ This could imply that the observed discrepancies between the LF-derived T2 and the experimental values might not be caused by an inaccuracy of the CALVADOS model. One could indeed hypothesize that a part of the T2 signal does arise from local backbone flexibility as probed by LFs, but another part could be due to local modifications of the chemical environment of residues due to transient contacts, and this is not probed by LFs. This could also explain the higher discrepancy between the LF-derived T2 values and the experimental ones caused by the introduction of phosphorylations (Figure 6), as these PTMs constitute a major chemical disruption in the local environment of the IDP.

### LFs, LCs and ***d_W LC_*** offer new insight into the local dynamics of IDPs

The similar behaviors of LFs and LCs for the tau mutant led us to investigate whether there was a local correlation between flexibility and curvature and if this correlation was homogeneous along the sequence. We computed the correlation between LCs and LFs along the sequence with a 30-residue-long gliding window (Figure SI-8). The correlation is very high (*r* > 0.85) and barely fluctuates for the repeat domains. It is comprised between 0.7 and 0.8 for the N-terminal regions, and even drops to 0.6 for the N1 domain. This inhomogeneity in the correlation profile between curvature and flexibility is difficult to interpret, but could be related to the functional role of the repeat domains. For the tau mutant, the correlation between LCs and smoothed LFs degrades for domains N-ter to P1. These domains also exhibit a closer proximity to other domains than prescribed by the WLC. The same pattern is observed for *α*-synuclein on a reduced scale (see discussion in SI and Figure SI-9). We can therefore ponder whether a high correlation between flexibility and curvature could have functional implications for the domains of IDPs, and on the other hand if a decorrelation could be symptomatic of more compact regions.

### Functional considerations

In 2021, Brotzakis et al. characterized weak and strong binding regions of tau repeats to MTs through MD simulations combined with experimental restraints.^55^ The weak binding regions correspond to the C-terminal part of the repeats, while the strong binding regions are centered on the SKXGS motif. These regions coincide with the pattern in flexibility and average curvature identified with LFs and LCs for the repeat domains (Figure 5), with the weak binding regions presenting less curvature and more rigidity than the strong binding regions. The strong binding regions interact with the MT surface at the interface between *α*- and *β*- tubulin monomers, and tend to bury between them, while the weak binding regions interact transiently with the monomer core. This was also shown to be the case for simulations of the repeat R2 alone with tubulins, in MD simulations without restraints.^88^ This binding mechanism of the repeats leads us to hypothesize that perhaps their flexibility and curvature profile in a free tau monomer is not a coincidence. It would indeed be easier for the strong binding regions to bury themselves at the interface if it was more curved and more flexible than the rest of the repeats. At the STO level, the more extended conformation on average of the repeat domains with regards to each other (Figure 3) would also favor the spreading of the repeats on the MT surface.

Another interesting finding is the lack of difference between the dynamic behavior of the pseudo-repeat R’ and of the repeats R1-4. R’ is often considered as the ’*ugly duckling*’ in tau studies due to its smaller homology with the other repeats. But a recent work by El Mammeri et al. shows that this pseudo-repeat might be of great relevance to tau functionality as its binding to MT is more favorable than the other repeats.^89^ The proximity of the C-ter domain might be a first lead in understanding why this is the case (Figure 3).

There might therefore be functional information encoded in the local dynamics and the STO of tau and of other IDPs, and the characterization proposed in this study might help unravel it further.

### Refining the paperclip model and impact of phosphorylating the AT8 epitope

An experimental characterization of the STO of the tau protein remains a challenge due to its disordered nature. A widely accepted model is the “paperclip” model that was first proposed by Jeganathan et al. in 2006. ^50^ It prescribes that the tau monomer adopts a paperclip-like conformation with the N-terminal and C-terminal regions staying at close distance to one another (FRET-derived distance of 23Å) even though the protein remains fully disordered. The C-ter domain is also measured to be at a FRET distance of 23Å from the R3 repeat domain (distance between residues 322 and 432), while the N-ter domain remains beyond the FRET distance from the R2 and R3 repeat domains (residues 17/18 and 291/322). Although the average distances measured in our study are not in quantitative agreement with those measurements, the overall behavior reported in Figure 3B is in full qualitative agreement. We do observe a closer proximity between the N-ter and C-ter domains than prescribed by the WLC model, and the C-ter domain is indeed closer to the repeats. Additionally, the N-ter domain is indeed closer to the repeats as it follows from the strong proximity between N2 and P1, but still remains further from R2 and R3 than the FRET distance would allow for a detection (the residues of the N-ter are further than 60Å on average from those of R2, while the FRET experiment from Jeganathan et al. would not have allowed accurate detection further than around 40 Å).

In a follow-up study two years later, Jeganathan et al. attempted to characterize the effect of different phosphorylation patterns by designing phosphomimetic mutants of tau with glutamic acid mutations.^90^ One of their mutants had pseudophosphorylations at sites 199, 202 and 205, very similar to our phosphorylation state. Their main finding was a drastic drop in measured FRET energy from the N-ter/C-ter interaction when the pseudophosphorylations were introduced, which corresponds to an increase of the N-ter/C-ter distance. The N-ter/repeat FRET interaction was already too weak to be accurately measured for the unphosphorylated tau mutant, but pseudophosphorylations still led to a lower FRET energy. These results are in agreement with our observations upon phosphorylations at S202 and T205. This leads us to combine the information obtained from the deviation coefficient map and the LFs and LCs profile to propose a refined paperclip model incorporating both intra- and interdomain relationships (Figure 8).

**Figure 8:**
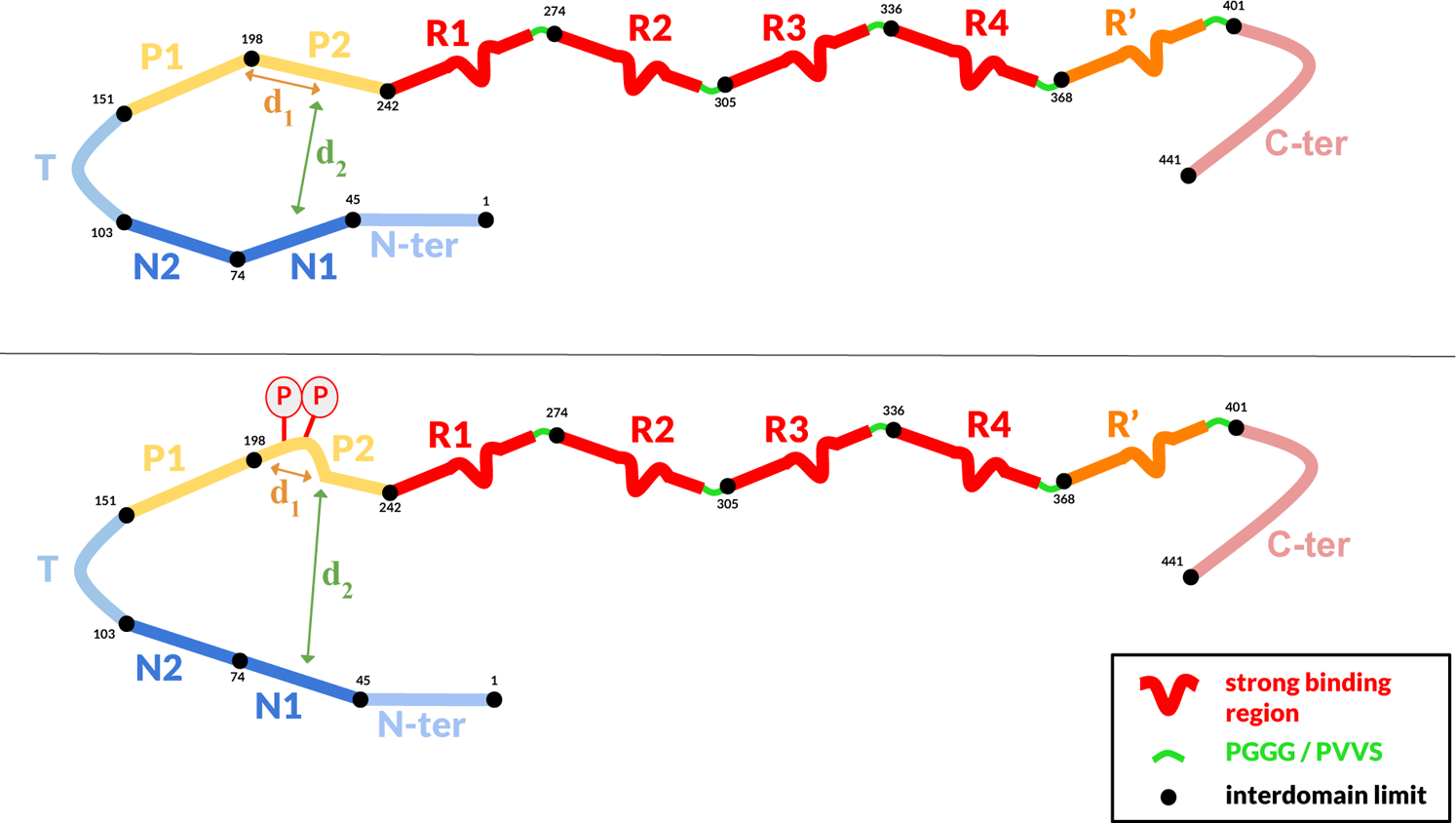
Statistical Tertiary Organization (STO) of the tau protein revealing the refined paperclip model. Upper panel: monomer organization without phosphorylations. Lower panel: monomer organization with phosphorylations at S202 and T205. The strong binding regions of the repeats are represented as curved following the LC profiles of Figure 4. *d*_1_ and *d*_2_ are the distances S198-T220 and G60-T220 respectively.

The STO of tau can be assumed to be relatively extended from domains P1 to R’. Domains N-ter, N1 and N2 are closer to P1 and P2 than the WLC model would prescribe, meaning that the domain spanning residues 103 to 150 would constitute a turn of sort. For this reason we decided to name it the Turn domain or T domain for short. The C-ter domain is closer to repeat domains and the increase in proximity compared to the WLC model could give rise to the FRET signals measured by Jeganathan et al. The resulting STO does resemble a paperclip, and the N-terminal “arm” is opened by AT8 phosphorylations. As far as we could tell, individual tau conformations however do not resemble proper paperclips, only their average behavior, translated by the STO, ressembles it. The STO should therefore be interpreted as a sort of ’archetypal configuration’, but must not be interpreted as a preferred physical conformation adopted by tau.

### Designing possible FRET and PRE/PRI experiments

In order to obtain FRET distances or PRE/PRI profiles, experimentalists must engineer protein mutants so as to strategically place cystein residues on which to graft their probes. This process is costly and it can be difficult to decide where to place the probe without a priori knowledge of the protein dynamics. Hence we decided to study more thoroughly two distances from our simulations in order to perhaps help guide future experimental characterization of the tau protein. We define the distances *d*_1_ and *d*_2_ as the distances between residues S198-T220 and G60-T220. Residues G60 and T220 can be considered as the central residues of the sequences of domains N1 and P2 respectively, and residue S198 is the first residue of the sequence of domain P2. We can imagine that the probe could be grafted at T220 by mutating it to cystein, while the two other residues could be mutated to tryptophans. The *d*_1_ distance could be used to probe the intra-domain compaction revealed in Figure 3B by phosphorylations, while the distance *d*_2_ would reveal whether the phosphorylations do repel the N1 domain from the P2 domain. We show the distributions of *d*_1_ and *d*_2_ and the distribution of the associated deviation coefficients for the monomeric simulation in Figure SI-10.

The distribution of *d*_1_ is slightly shifted towards lower values upon phosphorylations, while the distribution of *d*_2_ is clearly shifted towards higher values. Considering the deviation coefficients helps to visualize that the shift induced by phosphorylations is significant for both distances. These two distances could therefore be good observables to consider for further FRET and PRI experiments, and using the same mutant to obtain a PRE profile near residue T220 would also grant some visibility on the intradomain impact of phosphorylations. These experiments would help dissipate the remaining uncertainty on the exact mechanism by which phosphorylations impact the paperclip model and could then be easily compared with the CALVADOS ensembles.

## CONCLUSION

We used coarse-grain MD simulations with the CALVADOS model to investigate the conformational landscape of the disordered tau protein without and with phosphorylations. Characterizing the local compactness of an IDP as a deviation from a homogeneous wormlike chain allows us to highlight that disorder comes in various flavors, as recently observed by Chakraborty et al., ^36^ as we can define different domains along the tau sequence. The spatial repartition of these domains can be interpreted physically based on the average relative positions of their components, which we define as the Statistical Tertiary Organization (STO) of IDPs. We also used IDP-specific metrics to characterize the local curvature and flexibility of tau, and showed how the LFs are related with the protein dynamics and binding properties, even without the explicit modeling in the simulations of an interaction partner of tau. Finally, we combined results of both global and local scales to rediscuss the original paperclip model of tau and how it is impacted by phosphorylations. The resulting changes in the protein intra- and interdomain interaction pattern allow us to propose experimental setups that would help test our hypothesis.

## DATA AVAILABILITY

Experimental chemical shifts, T2 relaxation times, and all simulations described in this manuscript are available in the following Zenodo repository: https://zenodo.org/records/15979434

## Acknowledgement

The authors thank Dr. Alessia Lasorsa and Dr. Isabelle Landrieu for sharing the experimental NMR data; Dr. Caroline Smet-Nocca for fruitful discussions on NMR and on the results; Dr. Kresten Lindorff-Larsen and Dr. Giulio Tesei for fruitful discussions on the CALVADOS forcefield and on secondary chemical shifts. This work was supported by the ANR (MAGNETAU-ANR-21-CE29-0024) and by the “Initiative d’Excellence” program from the French State (Grant “DYNAMO”, ANR-11-LABX-0011-01 and grant “CACSICE”, ANR-11-EQPX-0008). Financial support from the IR INFRANALYTICS FR2054 for conducting the NMR research is gratefully acknowledged.

## Supporting Information Available

The following files are available free of charge.

- Supplementary_informations.pdf: contains the 9 figures listed as supplementary information in the paper, the table of the localCIDER results on the tau sequence, a discussion on the observed discrepancies between LF-derived T2 and experimental T2, and the application of LCs, LFs and *d_W LC_* to *α*-synuclein.

