## Supplementary information for "Across (conformational) space and (relaxation) time: using coarse-grain simulations to probe the intra- and interdomain dynamics of the tau protein"

Table SI-1: Results from the localCIDER[1] analysis of the Tau sequence. $f_+$ and $f_-$ are the fractions of positively and negatively charged residues respectively. The Fraction of Charged Residues (FCR) is defined as $FCR = f_+ + f_-$. The Net Charge Per Residue (NCPR) is defined as $FCR = f_+ - f_-$. $\kappa$ quantifies the patterning of oppositely charged residues and is defined in Ref.[2]

| | **Length** | $f_+$ | $f_-$ | **FCR** | **NCPR** | $\kappa$ |
|---|---|---|---|---|---|---|
| **Full sequence** | 441 | 0.13 | 0.13 | 0.26 | 0.00 | 0.18 |
| **Projection region** | 150 | 0.09 | 0.23 | 0.32 | -0.13 | 0.20 |
| **Proline Rich region (PRR)** | 90 | 0.17 | 0.03 | 0.20 | 0.13 | 0.11 |
| **Repeats region** | 159 | 0.17 | 0.09 | 0.26 | 0.08 | 0.10 |
| **C-ter region** | 41 | 0.05 | 0.12 | 0.17 | -0.07 | 0.34 |
| **Projection + PRR regions** | 241 | 0.12 | 0.15 | 0.27 | -0.03 | 0.22 |
| **R' + C-ter region** | 74 | 0.12 | 0.12 | 0.24 | 0.00 | 0.17 |

### Additional discussions

#### Discrepancies between the LF-derived T2 and the experimental T2

We computed the difference between the LFs-derived T2 and the experimental T2, then isolated the residue sequences for which the deviation was higher than the RMSE (Figure SI-SI-7). Excluding the termini, the sequences were 116-DEAAGHVTQA-125, 171-IPAKAPPAK-180, 213-PALPAPPT-220 and 309-VTKPVDLSKVT-319. All of them display values at least 0.05 s higher than the experimental measurement. This can be interpreted as backbone flexibility being too high for these fragments. The two sequences from the P1 and P2 domains are rich in prolines and display a similar succession of alanine residues. Since the CALVADOS forcefield does not integrate dihedral restrictions, one could wonder whether the prolines are still too flexible in the model as they would not capture cis/trans isomerization. Additionally, such sequences could be expected to display polyproline-II (PPII) helicity, since alanine, lysine and leucine have a strong PPII propensity near proline residues as well.[3] It is more difficult to rationalize why the two other sequences are not in agreement with the experimental data. On the other hand, residues in the N1 and C-ter domains display lower computed T2 values than the experimental measurements, which can be interpreted as backbone flexibility being too low. These domains contain a significant amount of negativaly charged residues such as glutamates, it can thus be hypothesized that the CALVADOS forcefield might slightly overestimate the electrostatic repulsions in these sections of the tau mutant.

#### Applications of LCs, LFs and $d_{WLC}$ to $\alpha$-synuclein

In order to probe whether LCs and LFs could provide valuable information for other IDPs, we performed a CALVADOS simulation of a monomer of $\alpha$-synuclein following the same protocol as for the Tau mutant. $\alpha$-synuclein is a 140-amino-acid-long IDP notably involved in Parkinson's disease.[4] It is composed of three domains : the N-terminal domain (residues 1-60), the

Non Amyloid Component (NAC, residues 61-95) and the C-terminal domain (residues 96-140). We computed the LCs, LFs and smoothed LFs (Figure SI-SI-8AB). Interestingly, LCs and LFs both observe a minimum at the residues located between the domains. Contrary to the repeats of Tau, this means that there is no highly flexible hinge between the domains but rather a more elongated and rigid part of the sequence, which could help separating said domains. We then assessed the correlation between LCs and the smoothed LFs with the same gliding window as for the Tau mutant (Figure SI-SI-8C). The local correlation is superior to 0.85 for most of domains NAC and C-ter, but drops to around 0.75 for the domain N-ter. In order to assess whether this could be related to the interdomain organization, we also computed the average distance map and the map of deviation coefficients $d_{WLC}$ (Figure SI-SI-8DE). The distance map reveals that the end of the C-ter domain is more extended than the other domains since the space between its isovalues is smaller. This is also characterized by the deviation coefficients, which range from 1.0 to more than 1.2 in most of the domain. On the other hand, residues of the NAC domain are slightly closer to one another than prescribed by the WLC model. Finally, the N-ter domain is ambivalent with deviation coefficients slightly lower than 1.0 for residues sequentially close, while all residues of the N-ter domain are not as close as prescribed from the N-terminal residues of the NAC domain. Note that the local compaction spots in Figure SI-9E concur with the contact maps obtained by Baul et al.[5] when using the SOP-IDP coarse-grain model, an de Bruyn et al.[6] when using all-atom MD simulations.

(2) Das, R. K.; Ruff, K. M.; Pappu, R. V. Relating sequence encoded information to form

and function of intrinsically disordered proteins. *Current Opinion in Structural Biology* **2015**, *32*, 102–112.

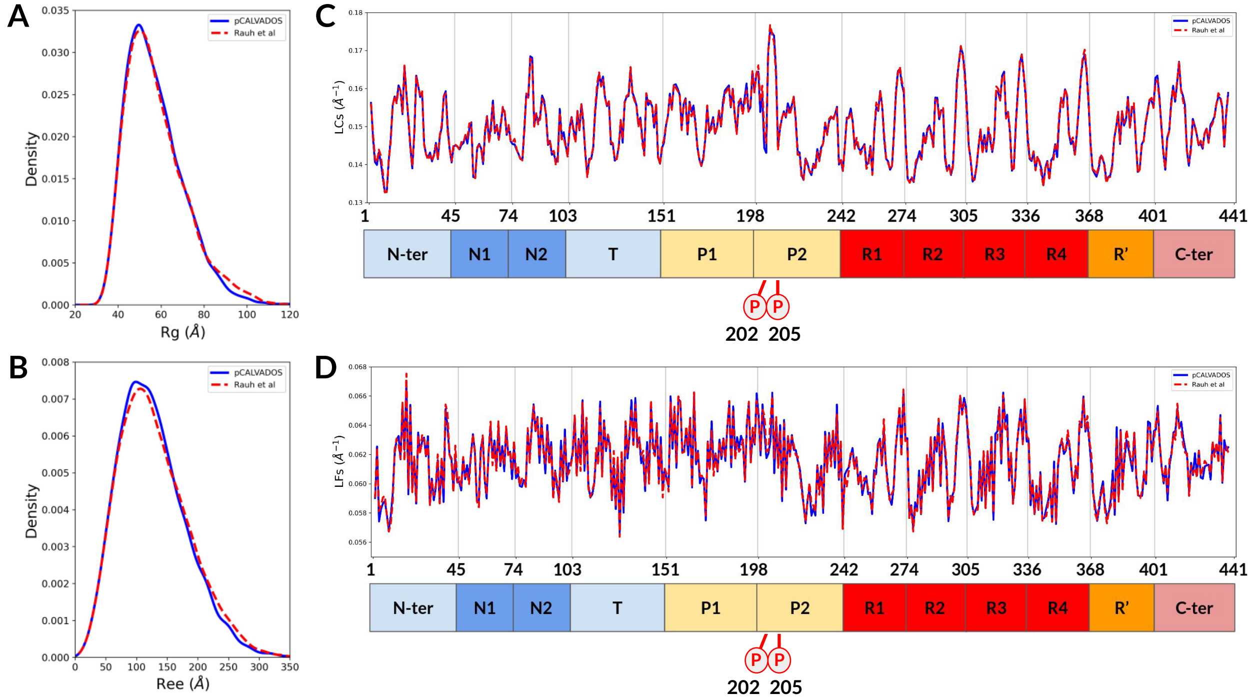


Figure SI-1: Comparison between the phosphorylated monomer simulations using the pCALVADOS parameters (in blue) or the parameters derived by Rauh et al[7] (in dotted red). A) Distribution of the radius of gyration Rg. B) Distribution of the end-to-end distance Ree. C) LCs. D) LFs.

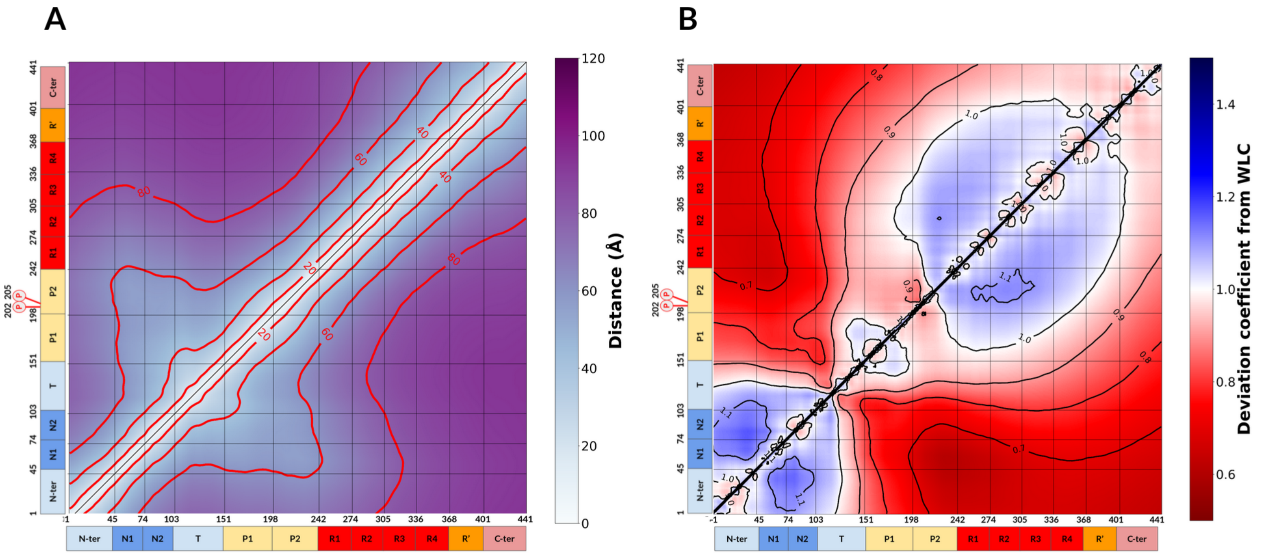


Figure SI-2: A) Average distances between residues in the unphosphorylated (lower part of the matrix) and ERK2-phosphorylated (upper part of the matrix) Tau mutant dimers. Increased distance is visualized by a deepening of the purple color. Isovalues 20Å, 40Å, 60Å, 80Å, 100Å and 120Å are plotted as red isolines. B) Deviation coefficient matrix from the WLC model. Deepening of the red color indicates a shorter average distance compared to the WLC model, deepening of the blue a longer average distance. Isovalues 0.6, 0.7, 0.8, 0.9, 1.0 and 1.1 are plotted as black isolines.

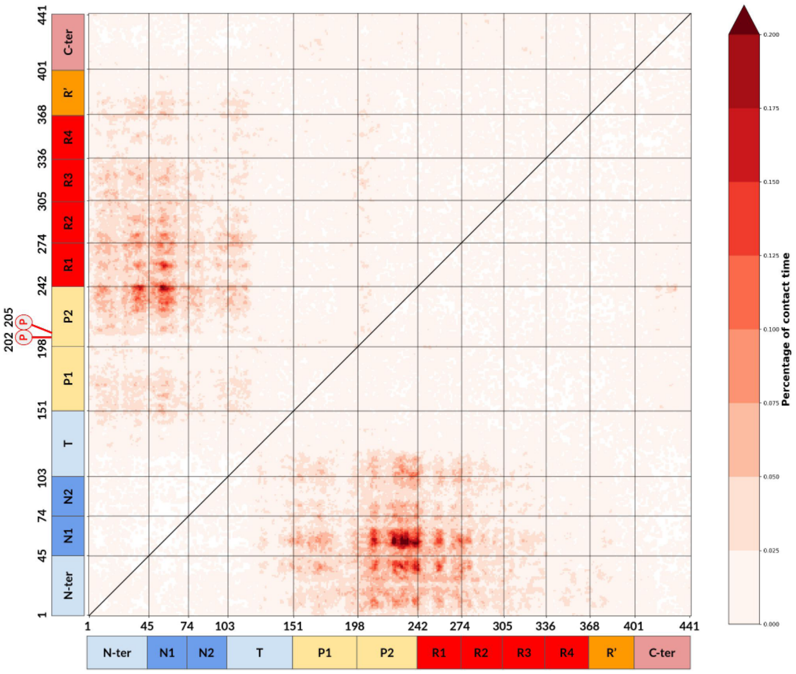


Figure SI-3: Contact map between monomers in the dimer simulations. Contacts between the unphosphorylated monomers (lower part of the matrix) and between the ERK2-phosphorylated monomers (upper part of the matrix). Increased contact time is visualized by a deepening of the red color.

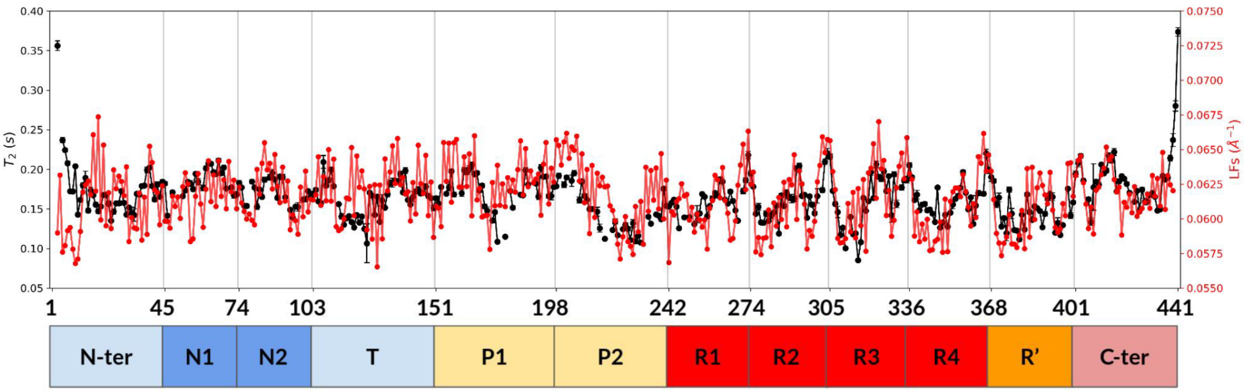


Figure SI-4: Manual superposition of the T2 relaxation time of the unphosphorylated Tau mutant (in black) and the LFs from the CALVADOS simulation (in red).

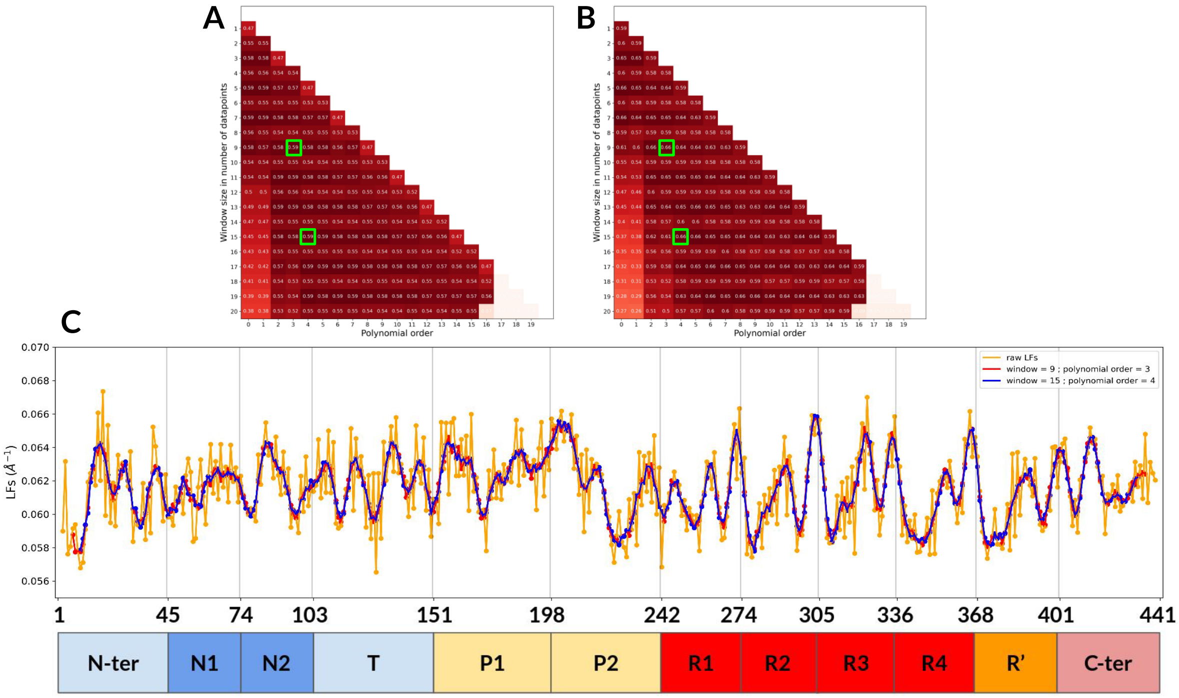


Figure SI-5: Correlation between the LFs and T2 when filtering LFs with a Savitzky-Golay filter at different window lengths and polynomial orders for A) the full sequence without the first and last 10 residues, B) the repeat domains. Maxima for low polynomial orders are reached in both cases in the green squares. C) Results of the filterings of LFs for a window of 9 residues and a polynomial order of 3 (in red), and a window of 15 residues and a polynomial order of 4 (in blue).

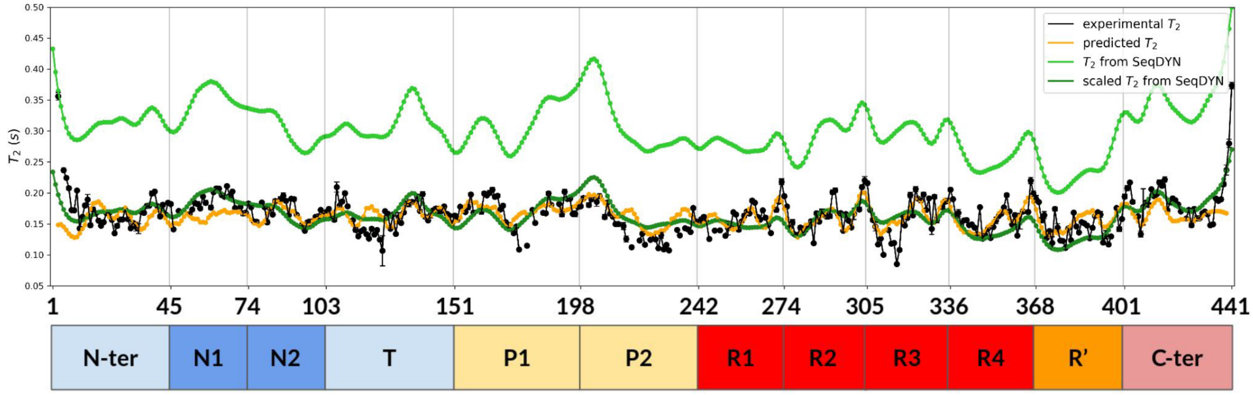


Figure SI-6: Comparison of the experimental T2 (in black), the LFs-derived T2 (in orange), the raw T2 from SeqDYN (pale green) and the scaled T2 from SeqDYN with a 0.54 scaling factor (in dark green).

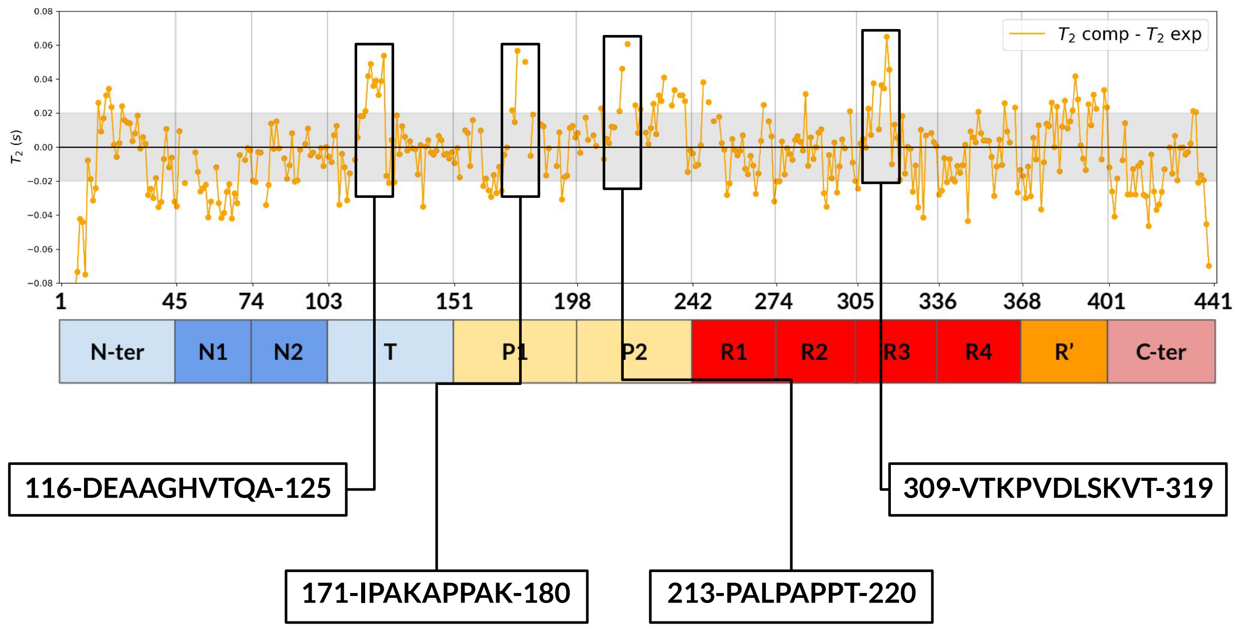


Figure SI-7: Difference between the predicted T2 from the smoothed LFs and the experimental T2. Positive values indicate that the predicted T2 is higher than the experimental T2. The RMSE value is 0.02s and is colored in grey. Sequences containing residues with a T2 difference higher than 0.04s are isolated with black squares.

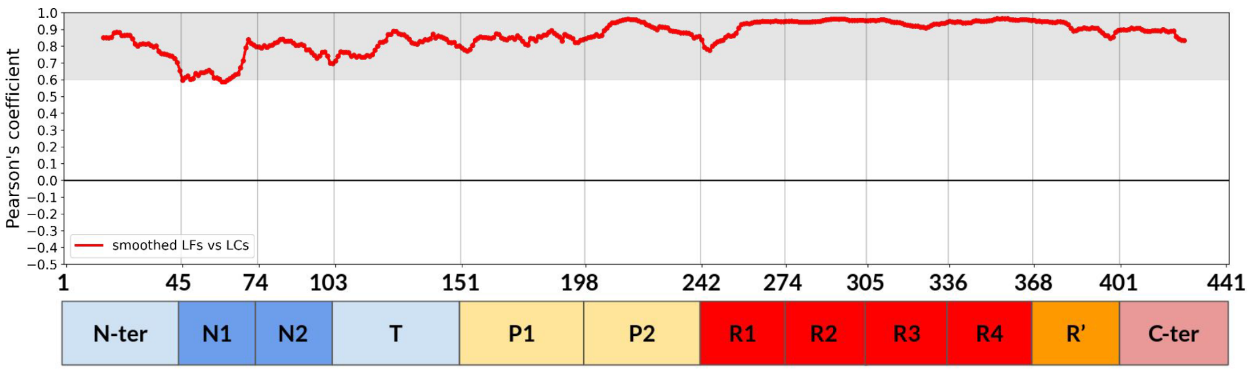


Figure SI-8: 30-residue gliding window of correlations between smoothed LFs and LCs. The value of the window is associated to the residue at the center of the considered sequence.

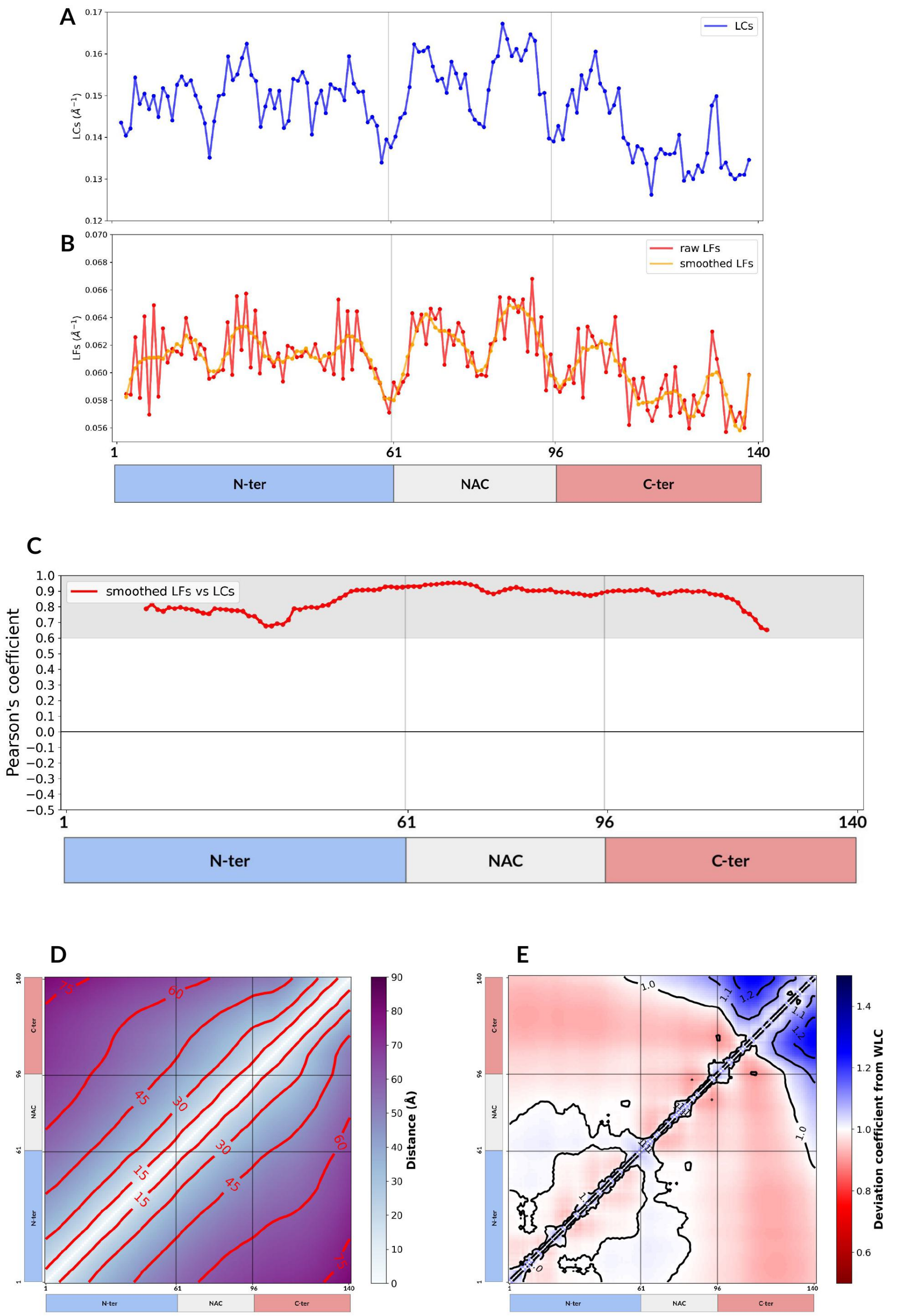


Figure SI-9: A) LCs calculated on the simulation of $\alpha$-synuclein. B) LFs (in red) and smoothed LFs (in orange), the smoothing was performed with a Savitzky-Golay filter with a window of 15 residues and a polynomial order of 4. C) 30-residue gliding window of correlations between smoothed LFs and LCs. The value of the window is associated to the residue at the center of the considered sequence. D) Average distances between residues (symetric matrix). Increased distance is visualized by a deepening of the purple color. Isovalues 15Å, 30Å, 45Å, 60Å and 75Å and 120Å are plotted as red isolines. E) Deviation coefficient matrix from the WLC model. Deepening of the red color indicates a shorter average distance compared to the WLC model, deepening of the blue a longer average distance. Isovalues 1.0 and 1.1 are plotted as black isolines.

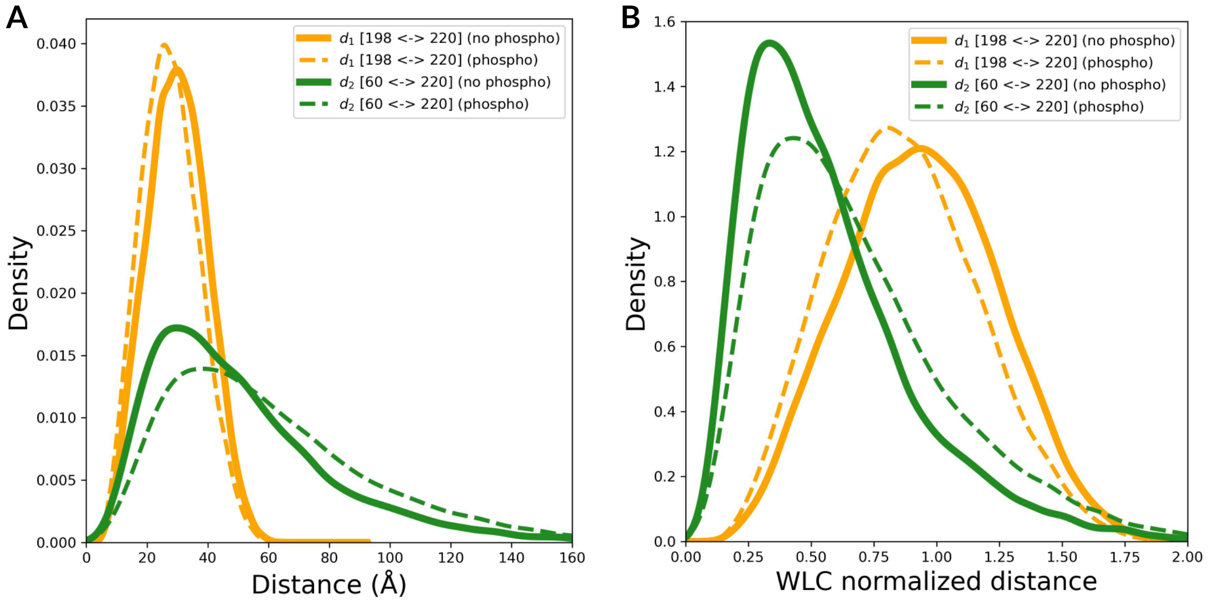


Figure SI-10: A) Distributions of the $d_1$ and $d_2$ distances (in orange and green respectively) corresponding to residues S198-T220 and G60-T220 respectively. Unphosphorylated case is in full line, phosphorylated case in dotted line. B) Distributions of the $d_1$ and $d_2$ distances normalized by their Worm-Like Chain model distance prediction.
